# Human GM-CSF/IL-3 enhance tumor immune infiltration in humanized HCC patient-derived xenografts

**DOI:** 10.1101/2023.10.05.561117

**Authors:** Kelley Weinfurtner, David Tischfield, George McClung, Jennifer Crainic, John Gordan, Jing Jiao, Emma E. Furth, Wuyan Li, Erena Tuzneen Supan, Gregory J. Nadolski, Stephen J. Hunt, David E. Kaplan, Terence P.F. Gade

## Abstract

**Background & Aims:** Responses to immunotherapies in hepatocellular carcinoma (HCC) are suboptimal with no biomarkers to guide patient selection. “Humanized” mice represent promising models to address this deficiency but are limited by variable chimerism and underdeveloped myeloid compartments. We hypothesized that expression of human GM-CSF and IL-3 increases tumor immune cell infiltration, especially myeloid-derived cells, in humanized HCC patient-derived xenografts (PDXs).

**Material and Methods:** NOG (NOD/Shi-*scid*/IL-2Rγ^null^) and NOG-EXL (huGM-CSF/huIL-3 NOG) mice conditioned with Busulfan underwent i.v. injection of human CD34+ cells. HCC PDX tumors were then implanted subcutaneously (SQ) or orthotopically (OT). Following serial blood sampling, mice were euthanized at defined tumor sizes. Tumor, blood, liver, and spleen were analyzed by flow cytometry and immunohistochemistry.

**Results:** Humanized NOG-EXL mice demonstrated earlier and increased human chimerism compared to humanized NOG mice (82.1% vs 43.8%, p<0.0001) with increased proportion of human monocytes (3.2% vs 1.1%, p=0.001) and neutrophils (0.8% vs 0.3%, p=0.02) in circulation. HCC tumors in humanized NOG-EXL mice had increased human immune cell infiltration (57.6% vs 30.2%, p=0.04), noting increased regulatory T cells (14.6% vs 6.8%, p=0.04), CD4+ PD-1 expression (84.7% vs 32.0%, p<0.01), macrophages (1.2% vs 0.6%, p=0.02), and neutrophils (0.5% vs 0.1%, p<0.0001). No differences were observed in tumor engraftment or growth latency in SQ tumors, but OT tumors required implantation at two rather than four weeks post-humanization for successful engraftment. Finally, utilizing adult bone marrow instead of fetal livers enabled partial HLA-matching to HCC tumors but required more CD34+ cells.

**Conclusions:** Human GM-CSF and IL-3 expression in humanized mice resulted in features more closely approximating the immune microenvironment of human disease, providing a promising model for investigating critical questions in immunotherapy for HCC.

**Impact and Implications:** This study introduces a unique mouse model at a critical point in the evolution of treatment paradigms for patients with hepatocellular carcinoma (HCC). Immunotherapies have become first line treatment for advanced HCC; however, response rates remain low with no clear predictors of response to guide patient selection. In this context, animal models that recapitulate human disease are greatly needed. Leveraging xenograft tumors derived from patients with advanced HCCs and a commercially available immunodeficient mouse strain that expresses human GM-CSF and IL-3, we demonstrate a novel but accessible approach for modeling the HCC tumor microenvironment.

## INTRODUCTION

Hepatocellular carcinoma (HCC) is a leading cause of cancer-related mortality worldwide, as the majority of patients present with unresectable disease and a median survival of less than two years^1,2^. Recently, immunotherapy has emerged as a first-line therapy for advanced HCC; however, these regimens remain suboptimal due to low response rates and high rates of adverse effects.^3,4^ Biomarkers of response to immunotherapy in other malignancies, including PD-1/PD-L1 expression level and tumor mutational burden, have not proven predictive of response in HCC.^3–7^ Early data suggest that immune cell phenotyping of HCC tumors may identify patients with greater likelihood of response to immunotherapy^8,9^; however, further understanding of the interaction between the immune microenvironment and immunotherapy outcomes is limited by the paucity of preclinical models that can recapitulate the heterogeneity of human disease.

While genetically engineered mouse models (GEMMs) have proven utility for the study of the impact of a single or combined genetic alteration(s) on tumor development and biology, they lack the heterogeneity and complexity of genetic and epigenetic alterations present in human HCC.^10,11^ Chemotoxic models allow for tumors to develop in a background of chronic liver disease with greater genetic diversity; however, tumorigenesis is more variable with longer latency and by a different mechanism than seen in patients.. Tumor formation in both of these syngeneic models occurs in immunocompetent mice, allowing for investigations into interactions between HCC tumors and the immune system with several important limitations. First, there are established differences in the targets of immunotherapy for human and murine homologs with several FDA-approved immunotherapies failing to bind to equivalent mouse targets.^12^ Second, GEMM tumors may have limited immunogenicity and often require additional strategies to mimic human immunosurveillance and immunotherapy responses.^13,14^ Lastly, while the murine immune system has frequently been used as a proxy for the human immune system, there are notable recognized, and likely unrecognized, differences in both innate and adaptive immunity.^15^

Patient-derived xenograft (PDX) models, in which human tumors are implanted into highly immunodeficient mice, overcome certain limitations of syngeneic mouse models based on their ability to recapitulate the heterogeneity of human tumors and patient responses to therapy which has led to their growing use in preclinical therapeutic trials.^16,17^ HCC PDXs have been described, but have largely been generated from patients with surgically cured, early-stage disease that likely represents less aggressive HCC variants that may never require systemic therapies. Recently, our lab and others have demonstrated the ability to generate HCC PDXs from biopsy samples, allowing for the development of PDX lines from patients with intermediate and advanced HCC – the patient population that will be considered for immunotherapy.^18,19^ However, the lack of immune system in these models has limited their utility for informing pressing questions in immuno-oncology.

To overcome this limitation, immunodeficient mice can be xenotransplanted with human hematopoietic cells to generate mice with a humanized immune systems (HIS).^20–22^ Severely immunodeficient mice, such as Nod-*scid* IL2Rγ^null^ (NSG) or NOD/Shi-*scid*/IL-2Rγ^null^ (NOG) mice, can be injected with human peripheral blood mononuclear cells (PBMCs) leading to good engraftment of memory and effector T cells; however, B cells and myeloid cells do not engraft well, and the study window is limited as these mice uniformly develop xenogeneic graft-versus-host disease (xGVHD) within 2 months of implantation.^23^ Alternatively, engraftment with human hematopoietic stem and progenitor cells (CD34+ cells) from fetal livers, cord blood, or bone marrow result in HIS mice that produce all lineages of hematopoietic cells, including both innate and adaptive immune cells, with delayed onset of xGVHD to over 6 months post-humanization.^24^ While myeloid progenitor cells engraft well in the bone marrow, there are low rates of mature myeloid cells in the periphery of these mice, including immune cell types important for tumor immune evasion such as macrophages, neutrophils, and dendritic cells.^22,25^ To date, the only published model of HCC in HIS mice demonstrated that the presence of the HCC PDX tumor led to significant decreases in all human cells in the peripheral blood relative to mice without implanted PDX, emphasizing the need for improved humanization strategies.^26^

The expression of various human cytokines has been shown to alter the proportion and functionality of different immune cell populations in HIS mice. Specifically, the expression of human granulocyte-macrophage colony-stimulating factor (GM-CSF) and interleukin-3 (IL-3) has been shown to increase not only overall hematopoietic cell engraftment, but also lead to a 3-fold increase in myeloid cells in the periphery.^27,28^ Interestingly, expression of these cytokines also leads to increased infiltration of regulatory T cells into all compartments, including tumor microenvironments.^27^ We aimed to investigate the influence of these human cytokines on HCC tumor growth and tumor immune infiltration in order to generate HIS HCC PDXs that more closely recapitulate human disease.

## MATERIALS AND METHODS

### Humanized mice

NOG (NOD.Cg-*Prkdc^scid^ Il2rg^tm1Sug^*) or NOG-EXL [NOD.Cg-*Prkdc^scid^ Il2rg^tm1Sug^* Tg(SV40/HTLV-IL3,CSF2)] mice were obtained from Taconic and frozen human fetal liver-derived CD34+ hematopoietic stem/progenitor cells were obtained from the University of Pennsylvania’s Stem Cell Xenograft Core. Only female mice were used due to prior data demonstrating significantly increased immune cell engraftment in female mice compare to male mice.^29^ The 6- to 8-week-old mice were conditioned with intravenous (i.v.) injection of 35 mg/kg busulfan and then injected with 300,000 human CD34+ cells by tail vein 24 hours later. Alternatively, frozen adult bone marrow (BM) CD34+ cells were obtained from Ossium Health to allow for partial HLA-matching to HCC PDX tumor prioritizing at minimum one allele match for *HLA-A*, HLA-B* and HLA-DRB1\**.

### Humanized HCC PDX generation

HCC PDXs were initially generated from clinical percutaneous biopsies of a human HCC tumors implanted into NSG mice and serially passaged as described previously.^19^ Tissue from these previously generated HCC PDX tumors was cut into approximately 1mm x 1mm pieces, frozen in freezing media (10% DMSO, 20% FBS, 70% DMEM), and stored in liquid nitrogen. After immune engraftment was confirmed by the presence of human CD45+ cells in the peripheral blood using flow cytometry at 4 weeks post-CD34+ cell transplant, the PDX tissue was thawed and rinsed twice with PBS. For subcutaneous (SQ) implants, the PDX tissue was aspirated into a 1 ML syringe containing 150 uL PBS and 150 uL Matrigel using an 18G needle and then injected SQ into the right flank of the humanized mice. For orthotopic implants, the humanized mice were anesthetized with isoflurane and a 1cm subcostal laparotomy incision was made to expose the liver. Using a No. 10 scalpel blade, a 3 mm incision was made on the surface of the exposed liver and a 1 mm x 1 mm piece of PDX tissue was placed into the incision and then covered with small drop of Matrigel (Corning, 354248). To ensure hemostasis, Surgicel (JNJ Medtech) was then applied over the tumor insertion site and the skin and peritoneum were closed in separate layers using absorbable sutures. Tumor growth was then monitored with calipers for SQ tumors or ultrasound for orthotopic tumors. Sham implantations with Matrigel only were also performed SQ or orthotopically. Mice were sacrificed once the tumor reached 20 mm in any direction or at end of experiment if no tumor growth. Tumors were considered engrafted if they reached a size of 4 x 4 mm in any dimension with increasing size on at least two consecutive measurements. Latency was defined as the number of days from implantation to tumor engraftment. Tumor doubling time was defined as number of days from tumor engraftment until volume of the tumor reached two times the volume of the tumor at engraftment.

### Immune cell analysis by flow cytometry

Immune engraftment was monitored by flow cytometry of peripheral blood samples obtained from retro-orbital blood collection every 2 weeks. At necropsy, peripheral blood, tumor, liver, and spleen were harvested. Tissue was mechanically dissociated through 70-micron filter and all samples underwent lysis of red blood cells with ACK lysing buffer (Quality Biological Inc) and blocking with Human TruStain FcX (Biolegend) prior to antibody staining using the antibodies listed in **Supplemental Table 1**. Flow cytometry panels were analyzed using the gating strategies shown in **Supplemental Figure 1**.

### Immunohistochemistry analysis

At necropsy, portions of tumor, liver, and spleen were fixed in formalin for subsequent paraffin embedding. After deparaffinization and rehydration of 4-μm-thick tissue sections, heat-induced epitope retrieval was performed in 10 mM sodium citrate buffer (pH 6.0). The slides were then blocked in StartingBlock T20 (Thermo Fisher Scientific) and then incubated with primary antibodies diluted in 5% bovine serum albumin in PBS with 0.2% Triton-X overnight. Primary antibodies used in this study are listed in **Supplemental Table 1**. Secondary antibodies conjugated to horseradish peroxidase (Vector Laboratories) were used and color was developed according to manufacturer protocols using DAB substrate kits counterstained with Mayer’s hematoxylin. Entire tissue sections were then imaged with a 20x objective using a Leica DM IRB microscope (Leica, Germany) with Leica imaging software. Cell counting was performed using ImageJ version 1.54h (National Institute of Health) in a blinded fashion by DT through analysis of high-power fields (20x) and averaging the number of cells within 3 separate areas each measuring 1 mm^2^.

### Cytokine/chemokine analysis

Plasma was obtained from biweekly peripheral blood samples collected as previously discussed. Cytokine and chemokine concentrations were determined using a custom Luminex assay (Bio-Techne) that included beads for CCL2, CCL4, CCL5, CCL20, CXCL1, CXCL5, CXCL10, CXCL13, granzyme A, GM-CSF, IL-2, IL-3, IL-4, IL-5, IL-6, IL-10, IL-12, IL-17/IL-17A, interferon-gamma, and TNF-alpha, as well as the following angiokines – angiopoietin-1, angiopoietin-2, Tie-2, and VEGF.

### Statistics

All data are expressed as mean ± standard deviation or standard error of mean. Differences between groups were assessed by either two-sided unpaired Student’s t-test or one-way analysis of variance (ANOVA) with Bonferroni post hoc test. Statistical analysis was performed on Prism 10 (GraphPad).

### Study approval

All mouse studies were approved by Institutional Animal Care and Use Committee at our institution. All animal experimental procedures were conducted in accordance with the approved protocols and Guide for the Care and Use of Laboratory Animals. Patient-derived xenografts were generated from patient biopsies in a prospective clinical study approved by University of Pennsylvania Institutional Review Board (IRB) #823696 and Corporal Michael J Crescenz VA Medical Center IRB #01779.

### Data availability

Individual FCS files for flow analysis and other primary data can be accessed from corresponding author on request.

## RESULTS

### Humanized NOG-EXL mice exhibit increased immune cell engraftment

HIS NOG-EXL mice demonstrated earlier and persistently increased huCD45+ peripheral blood cells compared to HIS NOG HCC mice with 12.1% vs 1.7% of cells at 4 weeks post-humanization (p<0.0001) and 82.1% vs 43.8% of cells at steady state (p<0.0001, **Figure 1A**).

**Figure 1:**
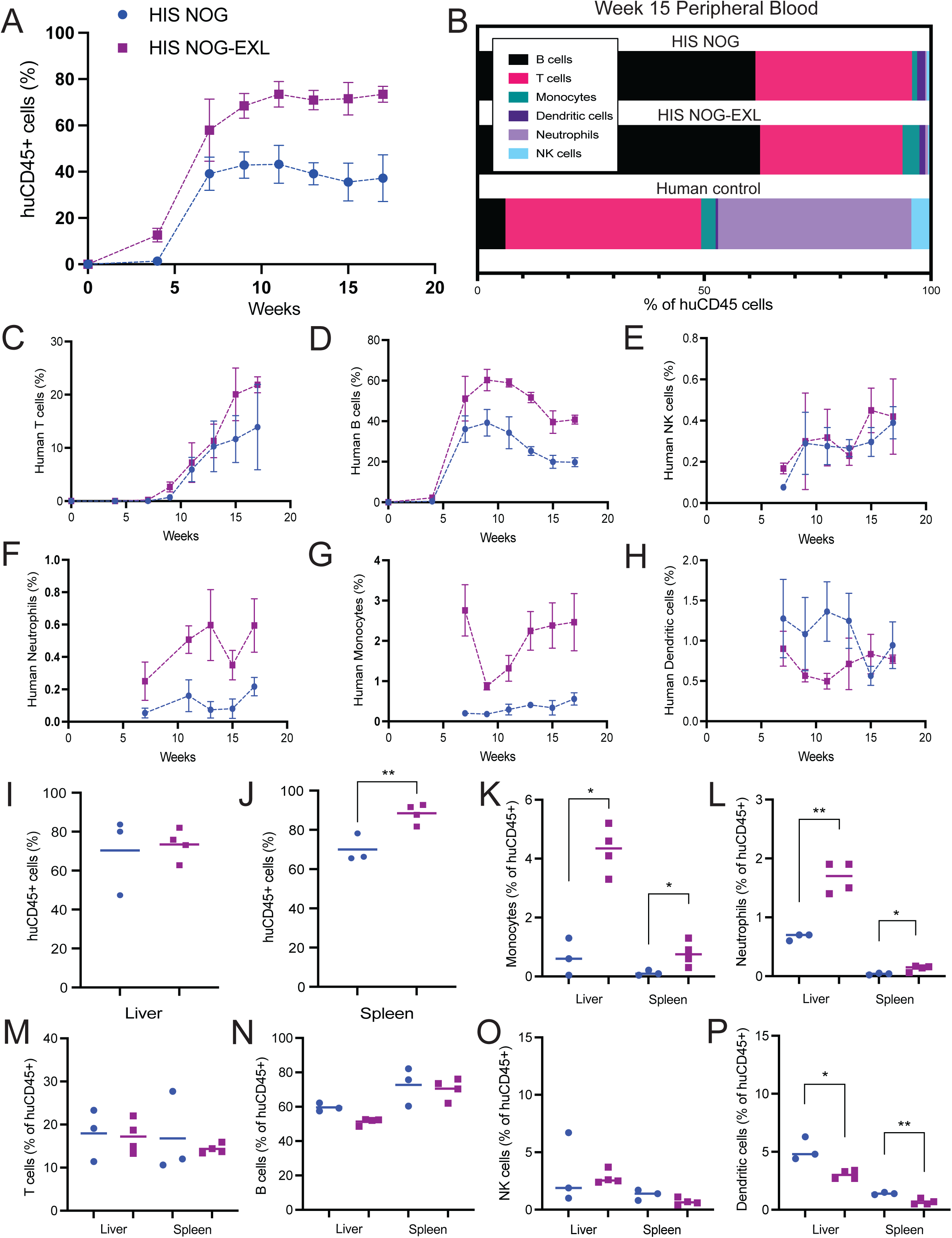
Impact of human GM-CSF and IL-3 on circulating human immune cells in HIS mice. (**A)** Percentage of peripheral blood cells that were human immune cells (huCD45+) by flow cytometry. (**B**) Percentage of human immune cells by subtype in peripheral blood at week 15 post-humanization. (**C-H**) Percentage of peripheral blood cells that were human immune cell subtypes from time of humanization. (**I-J**) Percentage of liver and spleen cells that were human immune cells (huCD45+). (**K-P**) Percentage of human immune cells by subtypes in liver and spleen. Blue circle = HIS NOG; Purple square = HIS NOG-EXL. *p<0.05; **p<0.01 (Student t-test).

All major immune cell types were represented in both groups, including B cells, T cells, NK cells, monocytes, dendritic cells, and neutrophils (**Figure 1B, Supp Figure 2**). HIS NOG-EXL mice showed a 2-fold increase in B-cells, a 2.7-fold increase in neutrophils, and a 5.5-fold increase in monocytes in the peripheral blood compared to NOG mice at 9 weeks post-humanization, although no difference in T-cells, NK cells, or dendritic cells was observed (**Figure 1C-H**). Human immune cell infiltration was also increased in the spleens (90.4% vs 73.2% of live cells, p=0.01) but not the livers (77.0 vs 70.4% of live cells, p=0.60) of HIS NOG-EXL mice relative to NOG mice (**Figure 1I-J**) with a proportional increase in monocytes (4.3% vs 0.6% of huCD45+ cells, p<0.01; 0.8% vs 0.1% of huCD45+ cells, p=0.049) and neutrophils (1.7% vs 0.7% of huCD45+ cells, p<0.01; 0.13% vs 0.04% of huCD45+ cells, p=0.03) in HIS NOG-EXL livers and spleens, respectively (**Figure 1K-P**). Notably, there were no differences in T-cell subtypes (CD4+, CD8+, Th1, Th17, Treg, naïve, effector) or T-cell PD-1 expression between the two cohorts in livers or spleens of the NOG-EXL and NOG HIS mice (**Supp Figure 3**).

**Figure 2:**
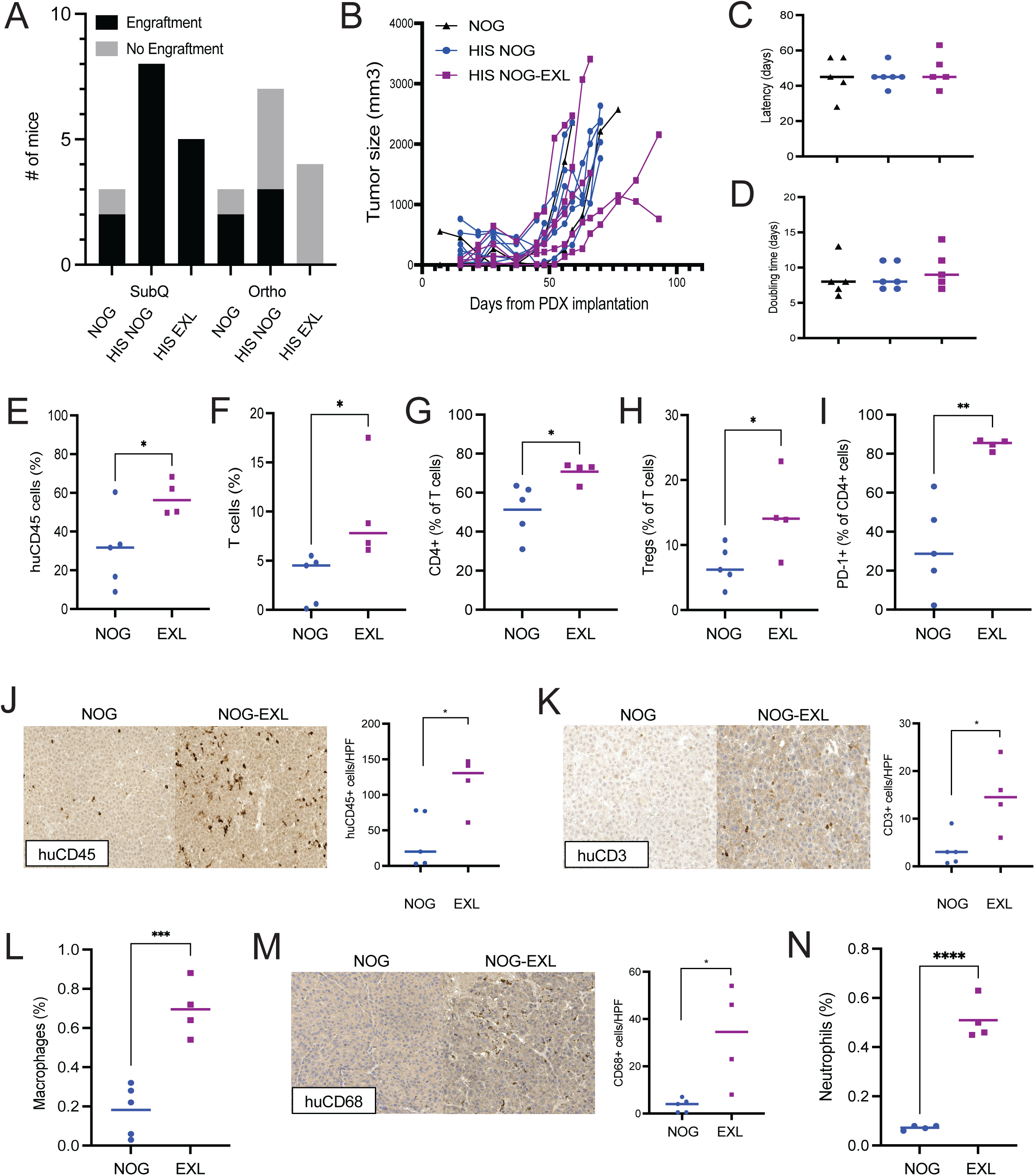
Influence of human GM-CSF and IL-3 on tumor immune cell infiltration in HIS mice. (A) Percentage of mice with tumor engraftment. (**B-D**) Tumor growth rate by individual tumor, latency, and doubling time; black triangle = NOG control; blue circle = HIS NOG; purple square = HIS NOG-EXL. (**E-F**) Percentage of tumor infiltrating immune cells that were human immune cells and human T cells by flow cytometry. (**G-I**) Percentage of tumor infiltrating human T cells that were CD4+, regulatory T cells, and expressing PD-1. (**J-K**) Number of human immune cells and human T cells per high power field by IHC. (**L**) Percentage of tumor infiltrating immune cells that are human macrophages by flow cytometry. (**M**) Number of human TAMs per high power field by IHC. (**N**) Percentage of tumor infiltrating immune cells that arehuman neutrophils by flow cytometry. Blue circle = HIS NOG; Purple square = HIS NOG-EXL. *p<0.05; **p<0.01, ***p<0.001, ****p<0.00001 (Student t-test).

**Figure 3:**
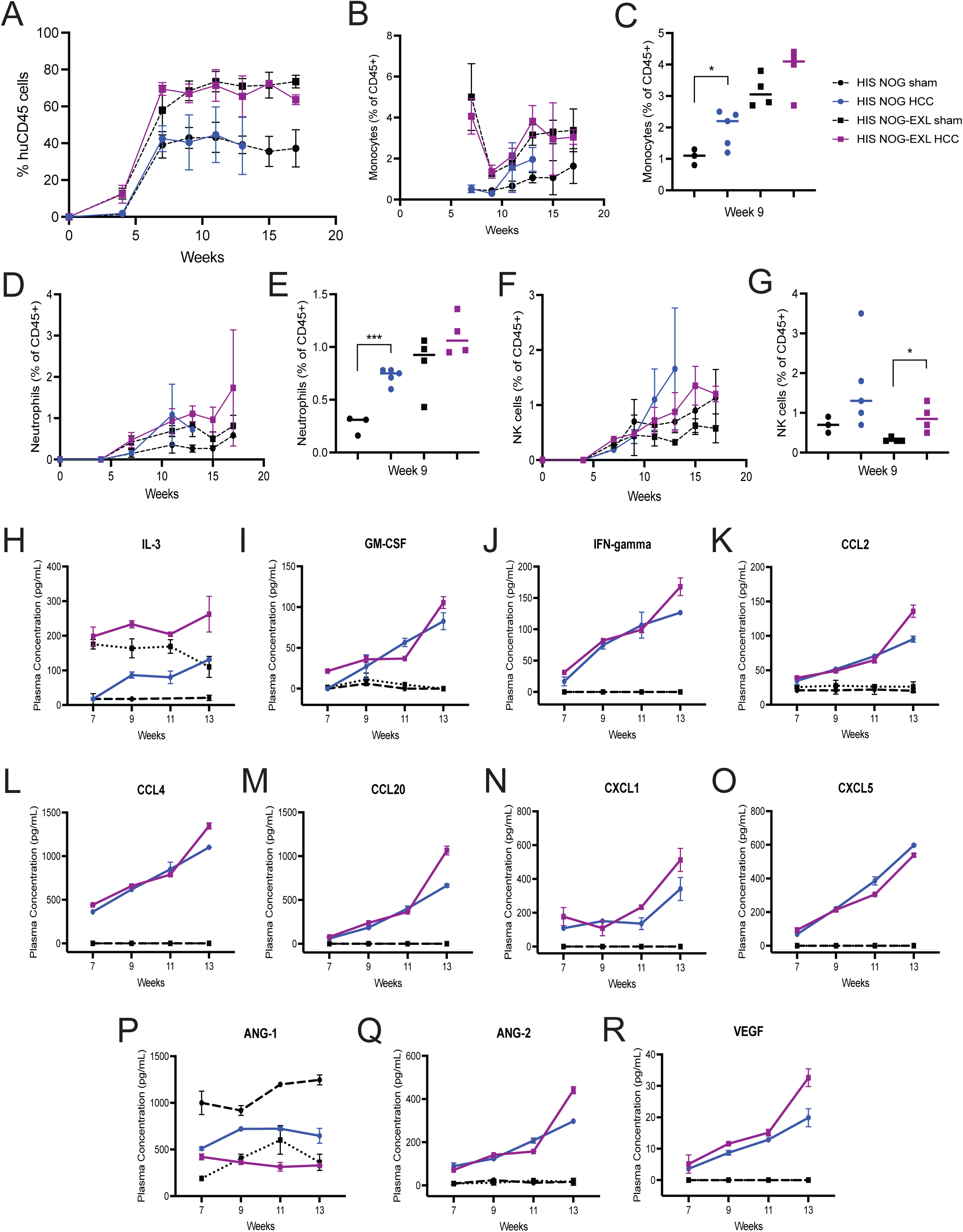
Effect of HCC tumor on circulating immune cells and chemokines in HIS mice. (**A**) Percentage of peripheral blood cells that were human immune cells (huCD45+) by flow cytometry in HIS mice with and without HCC PDX tumors. (**B-G**) Proportion of circulating human immune cells that were each subtype by flow cytometry in HIS mice with and without HCC PDX tumors. (**H-R**) Concentration of human cytokines and vascular growth factors in peripheral blood of HIS mice with and without HCC PDX tumors. Black circle, large dash = HIS NOG sham; Blue circle = HIS NOG HCC; Black square, small dash = HIS NOG-EXL sham; Purple square = HIS NOG-EXL. *p<0.05; ***p<0.001 (Student t-test).

### Humanization does not impact SQ HCC PDX tumor engraftment or growth rate

HIS mice were implanted SQ with human HCC PDX tumor (HCC #1), resulting in 100% engraftment regardless of mouse background (8/8 for HIS NOG, 5/5 for HIS NOG-EXL, **Figure 2A**). There was no difference in tumor growth between control NOG mice, HIS NOG mice, and HIS NOG-EXL mice as measured by growth latency (45.4, 45.5, and 48.4 days, respectively, p=0.63) or tumor doubling time (8.4, 8.7, and 9.8 days, respectively, p=0.63, **Figure 2B-D**). However, reduced tumor engraftment rates were observed for orthotopically implanted tumors in HIS mice – at 14 weeks post-tumor implantation, tumors engrafted in 3/8 (37.5%) HIS NOG mice and 0/4 (0%) HIS NOG-EXL mice compared to 2/3 (66.7%) NOG control mice (**Figure 2A**). Implanting PDX tumors orthotopically at 2 weeks rather than 4 weeks post-humanization improved engraftment significantly resulting in tumor engraftment in 11/13 (84.6%) HIS NOG-EXL mice at 15 weeks post-tumor implantation (**Supp Figure 4A**). These findings were confirmed in a second PDX line (HCC #2) implanted orthotopically two weeks after humanization with tumor engraftment in 7/13 (53.8%) HIS NOG-EXL mice compared to 2/3 (66.7%) in NOG-EXL controls at 22 weeks post-tumor implantation (**Supp Figure 4A**). Growth latency was specific to each PDX line; however, humanized mice had shorter growth latency than controls (**Supp Figure 4B**). Together, these results suggest an immune cell-mediated suppression of orthotopic tumor engraftment but enhanced growth rate that was not observed in the SQ compartment.

**Figure 4:**
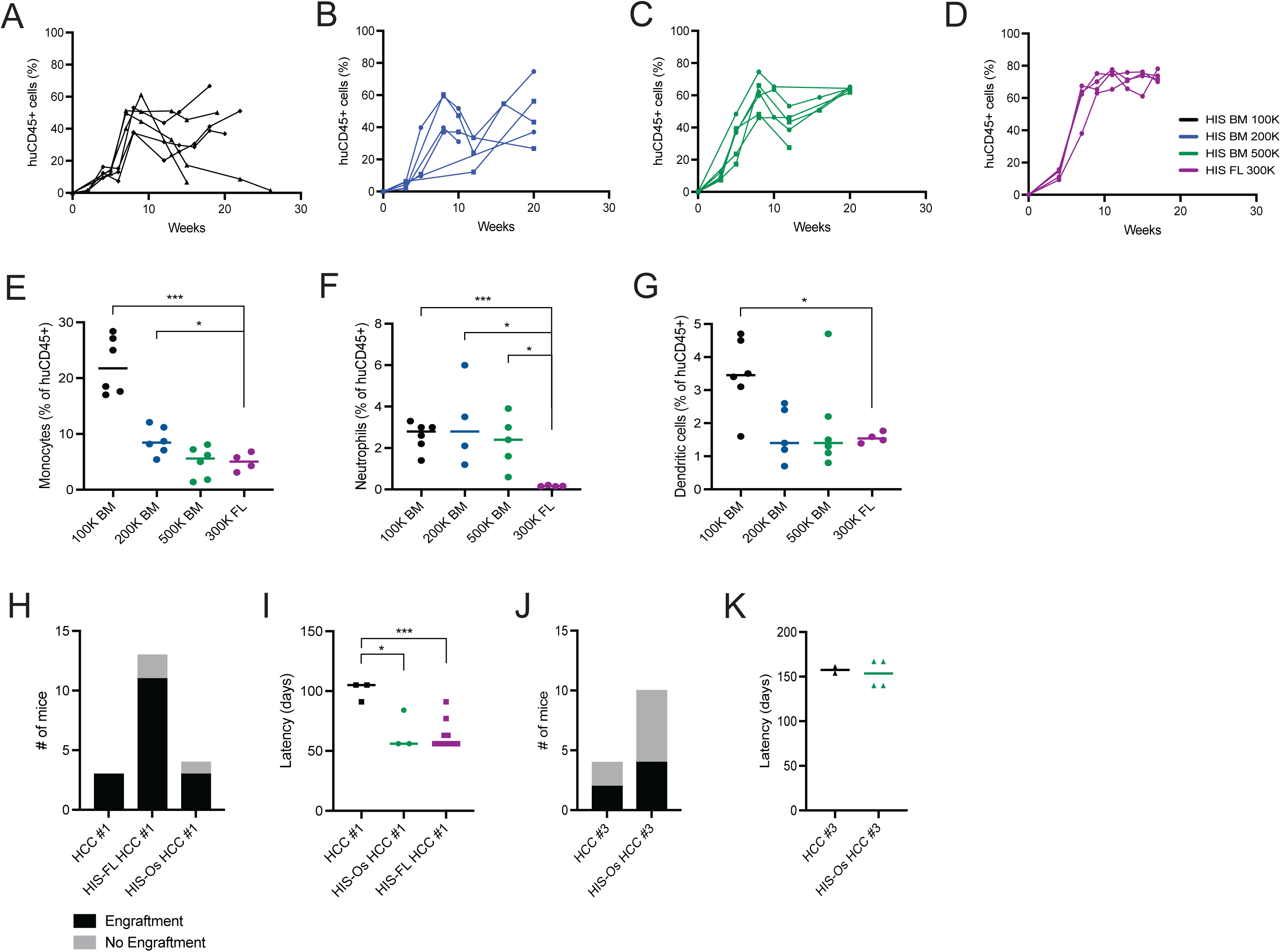
Immune engraftment in HIS mice depends on source of CD34+ cells. (**A-D**) Percentage of peripheral blood cells that were human immune cells (huCD45+) by flow cytometry for varying numbers of adult bone marrow derived CD34+ cells (BM-CD34+ cells) compared to fetal liver derived CD34+ cells (FL-CD34+ cells). (**E-G**) Proportion of human immune cells that were each subtypes in peripheral blood at week 7 post-humanization based on number and source of CD34+ cells. (**H-K**) NOG-EXL mice humanized with 500,000 BM-derived CD34+ (HIS-BM) or 300,000 FL-derived CD34+ (HIS-FL) were orthotopically implanted with human HCC PDX tumors two weeks after humanization. Tumor growth was monitored by ultrasound to determine tumor engraftment rates at 24 weeks and growth latency compared to tumors implanted in non-humanized NOG-EXL mice. *p<0.05; ***p<0.001 (Student t-test).

### Increased tumor immune infiltration in humanized NOG-EXL mice

At necropsy, HIS NOG-EXL mice demonstrated increased huCD45+ immune cell infiltration into SQ HCC tumors compared to HIS NOG mice by flow cytometry (57.6% vs 30.2% of live cells, p=0.04) and immunohistochemistry (117.3 vs 36.4 cells/HPF, p=0.02, **Figure 2E-K**). This included increased T-cell infiltration (flow cytometry: 23.1% vs 9.8% of live cells, p=0.04; IHC: 14.8 vs 3.3 cells/HPF, p=0.02) with proportional increases in CD4+ cells (70.8% vs 51.3% of T cells, p=0.03), CD4+ regulatory T cells (14.6% vs 6.8% of T cells, p=0.047), and CD4+ T cell PD-1 expression (84.7% vs 32.0% of CD4+ cells, p<0.01). HIS NOG-EXL mice also exhibited a proportional increase in tumor-associated macrophages (TAMs, flow cytometry: 1.2% vs 0.6% of CD45+ cells, p<0.01; IHC: 32.8 vs 3.4 cells/HPF, p=0.02, **Figure 2L-M**) and tumor-associated neutrophils (TANs, flow cytometry: 0.51% vs 0.07%, p<0.0001, **Figure 2N**).

### Presence of HCC tumor influences systemic cytokine and immune cell populations

The presence of HCC tumor did not impact overall peripheral blood chimerism in humanized mice (42.8% vs 40.5% in HIS NOG with HCC vs sham implanted mice, p=0.80; 68.4% vs 67.1% in HIS NOG-EXL HCC vs sham mice, p=0.71; **Figure 3A**); however, increased proportions of circulating monocytes (2.0% vs 1.1% in HIS NOG mice, p=0.047; 3.8% vs 3.2% in HIS NOG-EXL mice, p=0.19; **Figure 3B-C**), neutrophils (0.7% vs 0.3% in HIS NOG mice, p<0.01; 1.1% vs 0.8% in HIS NOG-EXL mice, p=0.16; **Figure 3D-E**), and NK cells (1.7% vs 0.7% in HIS NOG mice, p=0.20; 0.9% vs 0.3% in HIS NOG-EXL mice, p=0.02; **Figure 3F-G**) were observed in tumor bearing mice. When circulating chemokines were measured in the plasma of HIS mice, IL-3 concentration varied by mouse background with higher expression in HIS NOG-EXL mice regardless of HCC tumor presence; however, other chemokine levels increased as HCC tumors grew (**Figure 3H-O**), as did certain vascular growth factors (**Figure 3P-R**).

### Adult bone marrow CD34+ cells allow for partial HLA-matching but require greater number of cells to maintain adequate engraftment

Adult BM CD34+ cells are more readily available than fetal liver cells in larger quantities, allowing for partial matching by human leukocyte antigen (HLA)-type to HCC PDX tumor. Priority was given to matching at least one allele of *HLA-A*, HLA-B* and HLA-DRB1\**. When compared to using 300,000 fetal liver (FL)-derived CD34+, at least 500,000 adult BM-derived CD34+ cells were required to reach similar number of human immune cells in peripheral blood by week 9 post-humanization (42.0%, 49.1%, 59.4% vs 67.7% huCD45+ cells for 100,000, 200,000, 500,000 BM-derived CD34+ cells vs 300,000 FL-derived CD34+ cells, respectively, p<0.0001, **Figure 4A-D**). Even more notable, there was a significant decrease in circulating human immune cells after week 10 when using BM-derived CD34+ cells (28.2%, 25.8%, 41.8% vs 72.3% huCD45+ cells at week 12 for 100,000, 200,000, 500,000 BM-derived CD34+ cells vs 300,000 FL-derived CD34+ cells, respectively, p<0.0001). At least 500,000 BM-derived CD34+ were needed to consistently maintain chimerism past 14 weeks. Notably, the mice with decreasing chimerism using BM-derived CD34+ cells continued to display a significantly higher proportion of monocytes than FL-derived CD34+ control (17.7% vs 5.0% at week 9, p<0.0001). NOG-EXL mice humanized with 500,000 BM-derived CD34+ or 300,000 FL-derived CD34+ were implanted with HCC PDX tumors (HCC #1, #3) orthotopically in the liver two weeks after humanization. Tumor engraftment rate and growth latency were similar regardless of CD34+ cell source (**Figure 4H-K**).

## DISCUSSION

Here we present a novel humanized mouse model of HCC at a critical point in the evolution of treatment paradigms for patients with HCC. While combination immunotherapy strategies have become first line, response rates remain low with high rates of adverse events. Further, biomarkers of response to immunotherapy in other cancers have not been predictive of response in HCC, highlighting the unique tumor microenvironment in the liver and HCC tumors. In this context, animal models that recapitulate human HCC are greatly needed. Leveraging HCC PDX tumors derived from incurable HCCs, the population most likely to receive immunotherapy, and a commercially available immunodeficient mouse strain that expresses human GM-CSF and IL-3, we demonstrate a novel but accessible approach for modeling the HCC tumor microenvironment.

In this study, we demonstrate that HIS NOG-EXL mice have increased numbers of circulating human immune cells and increased human immune cell infiltration into HCC PDX tumors compared to standard immune deficient mouse strains (NOG, NSG). In addition to increased human myeloid cells in circulation and the tumor microenvironment, HIS NOG-EXL mice also showed an increased proportion of regulatory immune cells in the HCC tumor microenvironment, including Tregs and CD4+PD1+ T cells. Importantly, the model can be further optimized to allow for orthotopic rather than subcutaneous implantation of HCC PDX tumors, as well as partial HLA-matching of human hematopoietic cells to HCC tumor by using adult bone marrow-derived CD34+ cells instead of more limited fetal liver or cord blood. Taken together, these data represent an important advance in addressing the deficiency in animal models that can inform on pressing deficiencies in the treatment of HCC.

Humanized mouse models developed to date have been limited by variable and insufficient levels of human chimerism. In commercial models, 25% human chimerism of peripheral blood cells has been accepted as the cutoff for successful “humanization,” but this metric has not been optimized for investigations into the tumor immune microenvironment (TIME), especially “immune cold” tumors such as HCC.^30,31^ Recent data suggests that the degree of human immune infiltration into tumors in humanized models varies not only by patient and cancer type, but also based on the degree of HIS chimerism.^30^ In our HCC cohorts, the number of human immune cells in the TIME correlated with peripheral blood chimerism, in particular the number of peripheral human T cells, rather than splenic T cells as described previously. Importantly, HIS NOG-EXL mice showed an 87.4% increase in peripheral blood chimerism corresponding to a 90.7% increase in human immune cells in the TIME compared to HIS NOG mice. Collectively, these findings suggest that increased chimerism more effectively enables the study of immune subpopulations in the TIME and emphasize the importance of controlling for differences in chimerism in the interpretation of data acquired from humanized models. Importantly, increased chimerism did not result in earlier onset of xGVHD, as there was no evidence of xGVHD in these humanized mouse cohorts up to 22 weeks post-humanization.

Another major limitation of humanized models using standard immunodeficient mouse backgrounds is the underdeveloped myeloid compartment leading to low proportions of tumor infiltrating myeloid-derived cells, including MDSCs, TAMs, TANs, and CD103+DCs that all play an important role in tumor immune evasion.^21–22,25^ In addition to increasing overall chimerism and tumor infiltrating human immune cells, expression of human GM-CSF and IL-3 in HIS NOG-EXL mice also resulted in a greater than 3-fold increase in the proportion of monocytes/macrophages and neutrophils in the peripheral blood, liver, and HCC tumor. Interestingly, these mice also have an increased proportion of Tregs and PD-1+CD4+ T cells that was limited to the HCC tumor compartment though the mechanism for this increase remains unclear.

The humanization process applied to mice in this study did not impact subcutaneous HCC tumor engraftment, latency, or growth rate, but was associated with a significant decrease in orthotopic HCC engraftment. Orthotopic engraftment rates were improved by implanting PDX tumors at 2 rather than 4 weeks post-humanization, suggesting that the presence of larger numbers of human immune cells impairs tumor engraftment in the liver. Interestingly, we saw decrease latency of tumor formation in one of two HCC PDX lines implanted, suggesting the presence of immune cells may enhance tumor growth once tumors are engrafted. In the only other published model of humanized HCC PDXs, humanization of NSG mice increased growth rates of both SQ and orthotopic HCC tumors.^26,32^ This difference is unlikely to be due to mouse background, as NOG mice are functionally equivalent to NSG mice. Fetal livers were used as the source of human CD34+ hematopoietic stem/progenitor cells in both experiments, though Zhao et al. used CD34+ cells partially HLA-matched to HCC PDX tumors, suggesting the difference may relate to alloimmunity. However, in subsequent cohorts, the use of partially HLA-matched adult BM-derived CD34+ cells to the HCC PDX tumor did not result in changes in tumor growth rate when compared to mice humanized with FL-derived CD34+ cells. Tumor intrinsic factors may account for differences in the impact of humanized immune systems on tumor growth, as vascularity of tumors is variable and tumor-associated immune cells can have both pro-inflammatory and immunosuppressive phenotypes.

Despite these advantages, this study also emphasizes several important factors to consider when using this humanized PDX model. Increased myeloid cell differentiation results in impaired survival in NOG-EXL mice due to macrophage activation syndrome (MAS) that develops around 20 weeks post-humanization and has a similar clinical phenotype to hemophagocytic lymphohistiocytosis.^33,34^ The development of this phenotype is delayed and less pronounced in NOG-EXL mice than NSG-SGM3 [NOD.Cg-*Prkdc^scid^ Il2rg^tm1Wjl^* Tg(CMV-IL3,CSF2,KITLG)] mice, due to more physiologic levels of cytokine expression in NOG-EXL mice.^34,35^ Building on these data, novel immunodeficient mice that integrate human cytokines into mouse cytokine loci have been developed but are not yet commercially available.^36^ Another issue is the scarcity and cost of CD34+ cells, especially in enough quantity to allow for HLA-matching to PDX tumor in large cohorts of mice. While the frequency of long-term repopulating HSCs is highest in fetal livers, it is difficult to predict when tissue will be available and cost-prohibitive to bank large enough numbers for HLA-matching.^37,38^ The data presented here demonstrates that adult BM cells collected at autopsy from a commercial source are an alternative source of CD34+ cells that can be partially HLA-matched to HCC PDX tumors. Higher numbers of adult BM-derived CD34+ cells are required to obtain similar levels of immune engraftment, likely due to fewer long-term HSCs and increased progenitors undergoing terminal differentiation into myeloid cells in presence of human GM-CSF and IL-3.^39–41^ Various strains of mice are being developed to enhance the long-term engraftment of adult BM-derived or mobilized peripheral blood-derived CD34+ cells to allow for use of autologous CD34+ cells.^42–44^ Notably, this would require additional sample collection for HCC patients, even though not all of their tumors will develop into PDXs given ∼25% engraftment rates for HCC PDX’s.^18,19^

There are several notable limitations specific to this study. A single HCC PDX line was used in order to evaluate differences in humanized immune system attributable to the model rather than tumor intrinsic factors; however, the reproducibility of this model was subsequently demonstrated in two additional PDX lines. Future studies are planned to investigate differences between HCC PDX lines in this model system and how they compare to differences in the tumor immune microenvironment of parent biopsies. In addition, further investigation into the functional consequences of using adult bone marrow-derived versus fetal liver-derived CD34+ cells and the impact of partial HLA-typing on degree of alloimmune reaction are underway.

In summary, HIS HCC PDX models demonstrate multilineage immune infiltration in the peripheral blood, spleen, liver, and HCC tumor. Expression of human GM-CSF and IL-3 led to increased tumor infiltrating immune cells with a higher proportion of myeloid-derived and regulatory immune cells, suggesting NOG-EXL mice may be a more representative model for preclinical trials with immunotherapy than standard immunodeficient mice (NSG, NOG). Further study is needed to determine how this model recapitulates response to immunotherapy and autoimmune toxicity seen in patients.

## ABBREVIATIONS

BM: bone marrow
FL: fetal liver
GM-CSF: granulocyte-macrophage colony-stimulating factor
GEMMs: genetically engineered mouse models
HCC: hepatocellular carcinoma
HIS: humanized immune system
HLA: human leukocyte antigens
IL-3: interleukin-3
MAS: macrophage activation syndrome
MDSC: myeloid-derived suppressor cell
NK cells: natural killer cells
NOG: NOD.Cg-*Prkdc^scid^ Il2rg^tm1Sug^*
NOG-EXL: NOD.Cg- *Prkdc^scid^ Il2rg^tm1Sug^* Tg(SV40/HTLV-IL3,CSF2)
NSG: NOD scid gamma
NSG-SGM3: NOD.Cg- *Prkdc^scid^ Il2rg^tm1Wjl^*Tg(CMV-IL3,CSF2,KITLG)
PBMC: peripheral blood mononuclear cells
PD-1: programmed cell death protein 1
PD-L1: programmed cell death protein ligand 1
PDX: patient-derived xenograft
SQ: subcutaneous
TAM: tumor-associated macrophage
TAN: tumor-associated neutrophil
TIME: tumor immune microenvironment
Treg: regulatory T cell
xGVHD: xenogeneic graft-versus-host disease

## ACKNOWLEDGEMENTS

Research reported in this publication was supported by Penn Center for Precision Medicine Accelerator Grant, VA CSR&D I01 CX-001933, SIO-Astra Zeneca 2022 Research Grant, and DOD Translational Team Science Award CA220654P2. KW is supported by CTSA KL2 Mentored Career Development Award (5KL2TR001879-08). The authors would also like to acknowledge the Stem Cell Xenograft Core (RRID: SCR_010035), the Center for Molecular Studies in Digestive and Liver Diseases (P30DK050306), and the Molecular Pathology and Imaging Core (RRID: SCR_022420).

## Author Contributions

KW: designed research studies, conducted experiments, analyzed data, wrote the manuscript

DT, TPFG: designed research studies, conducted experiments, analyzed data, revised manuscript

GM, JC, JJ, WL: conducted experiments, analyzed data, revised manuscript JG, EEF: analyzed data, revised manuscript

GJN, SJH: designed research studies, revised manuscript

DEK: designed research studies, analyzed data, revised manuscript

## REFERENCES

1. Author names in bold designate shared co-first authorship

1. Bray F, Ferlay J, Soerjomataram I et al. Global cancer statistics 2018: GLOBOCAN estimates of incidence and mortality worldwide for 36 cancers in 185 countries. CA. Cancer J. Clin 2018; 68: 394–424.

2. Park JW, Chen M, Colombo M et al. Global patterns of hepatocellular carcinoma management from diagnosis to death: The BRIDGE Study. Liver Int. 2015;35:2155–2166.

3. Finn RS, Qin S, Ikeda M et al. Atezolizumab plus Bevacizumab in Unresectable Hepatocellular Carcinoma. N. Engl. J. Med. 2020;382:1894–1905.

4. Abou-Alfa GK, Lau G, Kudo M et al. Tremelimumab Plus Durvalumab in Unresectable Hepatocellular Carcinoma. NEJM Evidence. 2022;1(8):1–12.

5. Zhu AX, Abbas AR, Ruiz de Galarreta M et al. Molecular correlates of clinical response and resistance to atezolizumab in combination with bevacizumab in advanced hepatocellular carcinoma. Nature Medicine. 2022;28:1599–1611.

6. Pinter M, Jain RK, and Duda DG. The Current Landscape of Immune Checkpoint Blockade in Hepatocellular Carcinoma: A Review. JAMA Oncol. 2021;7(1):113–123.

7. Llovet JM, Montal R, Sia D, and Finn RS. Molecular therapies and precision medicine for hepatocellular carcinoma. Nat Rev Clin Oncol. 2018;15:599–616.

8. Sia D, Jian Y, Martinez-Quetglas I et al. Identification of an Immune-specific Class of Hepatocellular Carcinoma Based on Molecular Features. Gastro. 2017;153:812–826.

9. Montironi C, Castet F, Haber PK, et al. Inflamed and non-inflamed classes of HCC: a revised immunogenomic classification. Gut. 2023;72:129–140.

10. Brown ZJ, Heinrich B, and Greten TF. Mouse models of hepatocellular carcinoma: an overview and highlights for immunotherapy research. Nat Rev Gastro Hep. 2018;15:536–554.

11. Olson B, Li Y, Lin Y et al. Mouse models for cancer immunotherapy research. Cancer Discov. 2018; 1358–1365.

12. Magiera-Mularz K, Kocik J, Musielak B et al. Human and mouse PD-L1: similar molecular structure, but different druggability profiles. iScience. 2020;24(1):101960.

13. Ruiz de Galarreta M, Bresnahan E, Molina-Sanchez P et al. B-catenin activation promotes immune escape and resistance to anti-PD-1 in hepatocellular carcinoma. Cancer Discov. 2019;9(8):1125–1141.

14. Wang J, Perry CJ, Meeth K et al. UV-induced somatic mutations elicit a functional T cell response in the YUMMER1.7 mouse melanoma model. Pigment Cell Melanoma Res. 2017;30(4):428–435.

15. Mestas J and Hughes CCW. Of mice and not men: differences between mouse and human immunology. J. Immunol. 2004;172:2731–2738.

16. Gao H, Korn JM, Ferretti S et al. High-throughput screening using patient-derived tumor xenografts to predict clinical trial drug response. Nat Med. 2015;21:1318–25.

17. Byrne AT, Alferez DG, Amant F et al. Interrogating open issues in cancer precision medicine with patient-derived xenografts. Nat. Rev. Cancer. 2017;17(4):254–268.

18. Blumer T, Fofan I, Matter MS et al. Hepatocellular Carcinoma Xenografts Established From Needle Biopsies Preserve the Characteristics of the Originating Tumors. Hepatol. Commun. 2019;3:971–986.

19. Tischfield D, Ackerman D, Noji M et al. Establishment of Hepatocellular Carcinoma Patient-Derived Xenografts from Image-Guided Percutaneous Biopsies. Sci Rep. 2019;9:10546.

20. Yahata T, Ando K, Nakamura Y et al. Functional human T lymphocyte development from cord blood CD34+ cells in nonobese diabetic/Shi-scid, IL-2 receptor null mice. J. Immunol. 2002;169:204–209.

21. Shultz LD, Lyons BL, Burzenski LM et al. Human lymphoid and myeloid cell development in NOD/LtSz-scid IL2R gamma null mice engrafted with mobilized human hemopoietic stem cells. J Immunol. 2005;174:6477–89.

22. Shultz LD, Brehm MA, Garcia-Martinez JV, and Greiner DL. Humanized mice for immune system investigation: progress, promise and challenges. Nat Rev Immunol. 2012;12:786–98.

23. King MA, Covassin L, Brehm MA et al. Human peripheral blood leucocyte non-obese diabetic-severe combined immunodeficiency interleukin-2 receptor gamma chain gene mouse model of xenogeneic graft-versus-host-like disease and the role of host major histocompatibility complex. Clin Exp Immunol. 2009;157(1):104–118.

24. Sonntag K, Eckert F, Welker C et al. Chronic graft-versus-host-disease in CD34(+)-humanized NSG mice is associated with human susceptibility HLA haplotypes for autoimmune disease. J Autoimmun. 2015;62:55–66.

25. Tanaka S, Saito Y, Kunisawa J et al. Development of mature and functional human myeloid subsets in hematopoietic stem cell-engrafted NOD/SCID/IL2ryKO mice. J Immunol. 2012;188(12):6145–6155.

26. Zhao Y, Shuen TWH, Toh TB et al. Development of a new patient-derived xenograft humanised mouse model to study human-specific tumour microenvironment and immunotherapy. Gut. 2018;67:1845–1854.

27. Bilerbeck E, Barry WT, Mu K et al. Development of human CD4+FoxP3+ regulatory T cells in human stem cell factor-, granulocyte-macrophage colony-stimulating factor-, and interleukin-3-expressing NOD-SCID IL2R-gamma(null) humanized mice. Blood. 2011;117(11):3076–3086.

28. Maser IP, Hoves S, Bayer C et al. The tumor milieu promotes functional human tumor-resident plasmacytoid dendritic cells in humanized mouse models. Front Immunol. 2020:11;2082.

29. Notta F, Doulatoy S, and Dick J. Engraftment of human hematopoietic stem cells is more efficient in female NOD/SCID/IL-2Rgc-null recipients. Blood. 2010 May 6;115(18):3704–7.

30. Marin-Jimenez JA, Capasso A, Lewis MS et al. Testing cancer immunotherapy in a human immune system mouse model: Correlating treatment response to human chimerism, therapeutic variables, and immune cell phenotypes. Front Immunol. 2021;12:607282. Accessed 4/24/24.

31. huNOG-EXL Model Portfolio. Taconic Biosciences. https://www.taconic.com/products/mouse-rat/nog-portfolio/humanized-immune-system-mice/hunog-exl. Accessed 4/24/24.

32. Zhao Y, Wang J, Liu WN, et al. Analysis and Validation of Human Targets and Treatments Using a Hepatocellular Carcinoma Immune Humanized Mouse Model. Hepatol. 2021;74:1395–1410.

33. Tarrant JC, Binder ZA, Bugatti M et al. Pathology of macrophage activation syndrome in humanized NSGS mice. Res Vet Sci.2021;134:137–146.

34. Willis E, Verrelle J, Banerjee E et al. Humanization with CD34-positive hematopoetic stem cells in NOG-EXL mice results in improved long-term survival and less severe myeloid cell hyperactivation phenotype relative to NSG-SGM3 mice. Vet Pathol. 2024: 3009858231222216.

35. Comparison Guide: huNOG-EXL, NSG-SGM3, and MISTRG. Taconic Biosciences. https://www.taconic.com/resources/comparison-guide-hunog-exl-nsg-sgm3-and-mistrg. Accessed 4/24/24.

36. Rongvaux A, Willinger T, Martinek J et al. Development and function of human innate immune cells in a humanized mouse model. Nat Biotechnol. 2014;32(4):364–372.

37. Holyoake TL, Nicolini FE, and Eaves CJ. Functional differences between transplantable human hematopoietic stem cells from fetal liver, cord blood, and adult marrow. Exp Hematol. 1999;27(9):1418–1427.

38. Lepus CM, Gibson TF, Gerber SA et al. Comparision of human fetal liver, umbilical cord blood, and adult blood hematopoietic stem cell engraftment in NOD-*scid*/γc^−/−^, Balb/c-*Rag1*^−/−^γc^−/−^, and C.B-17-*scid*/bg immunodeficient mice. Hum Immunol. 2009;70(10):790–802.

39. Sippel TR, Radtke S, Olsen TM et al. Human hematopoietic stem cell maintenance and myeloid cell development in next-generation humanized mouse models. Blood Adv. 2019;3(3):268–274.

40. Hess NJ, Lindner PN, Vazquez J et al. Different Human Immune Lineage Compositions are Generated in Non-Conditioned NBSGW Mice Depending on HSPC Source. Front Immunol. 2020;11:573406.

41. Walcher L, Hilger N, Wege AK et al. Humanized mouse model: Hematopoietic stem cell transplantation and tracking using short tandem repeat technology. Immun Inflamm Dis. 2020:8(3):363–370.

42. Strowig T, Rongvaux A, Rathinam C et al. Transgenic expression of human signal regulatory protein alpha in Rag2^−/−−/−^ mice improves engraftment of human hematopoietic cells in humanized mice. PNAS. 2011;108(32):13218–13223.

43. Saito Y, Ellegast JM, Rafiei A et al. Peripheral blood CD34+ cells engraft in human cytokine knock-in mice. Blood. 2016;128(14):1829–1833.

44. Chiorazzi M, Martinek J, Krasnick B et al. Autologus humanized PDX modeling for immuno-oncology recapitulates features of the human tumor microenvironment. J Immunother Cancer. 2023;11(7):e006921.

