## Supplementary figures and images for "Human GM-CSF/IL-3 enhance tumor immune infiltration in humanized HCC patient-derived xenografts"

### Supplemental Figure 1

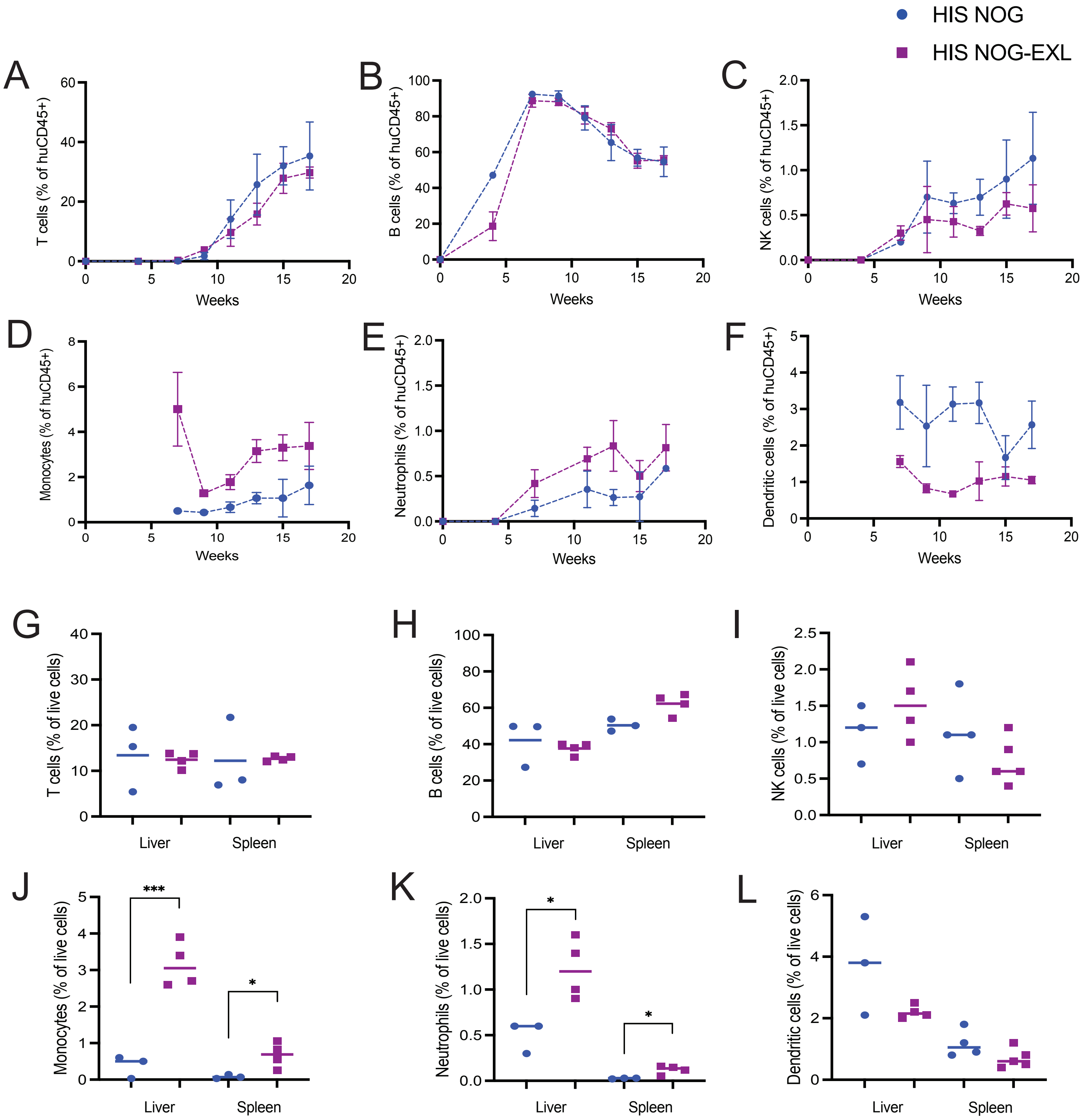

### Supplemental Figure 2

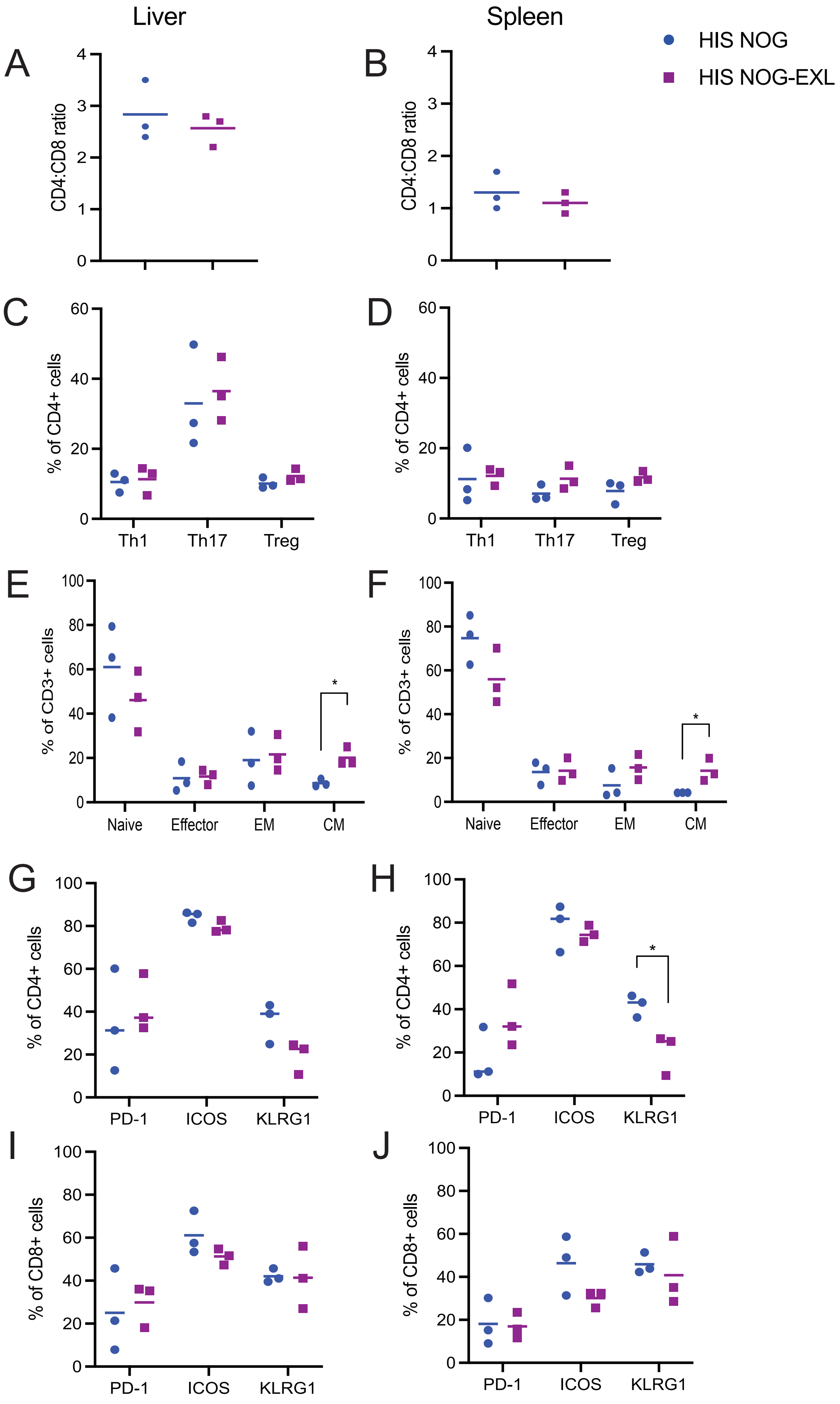

### Supplemental Figure 3

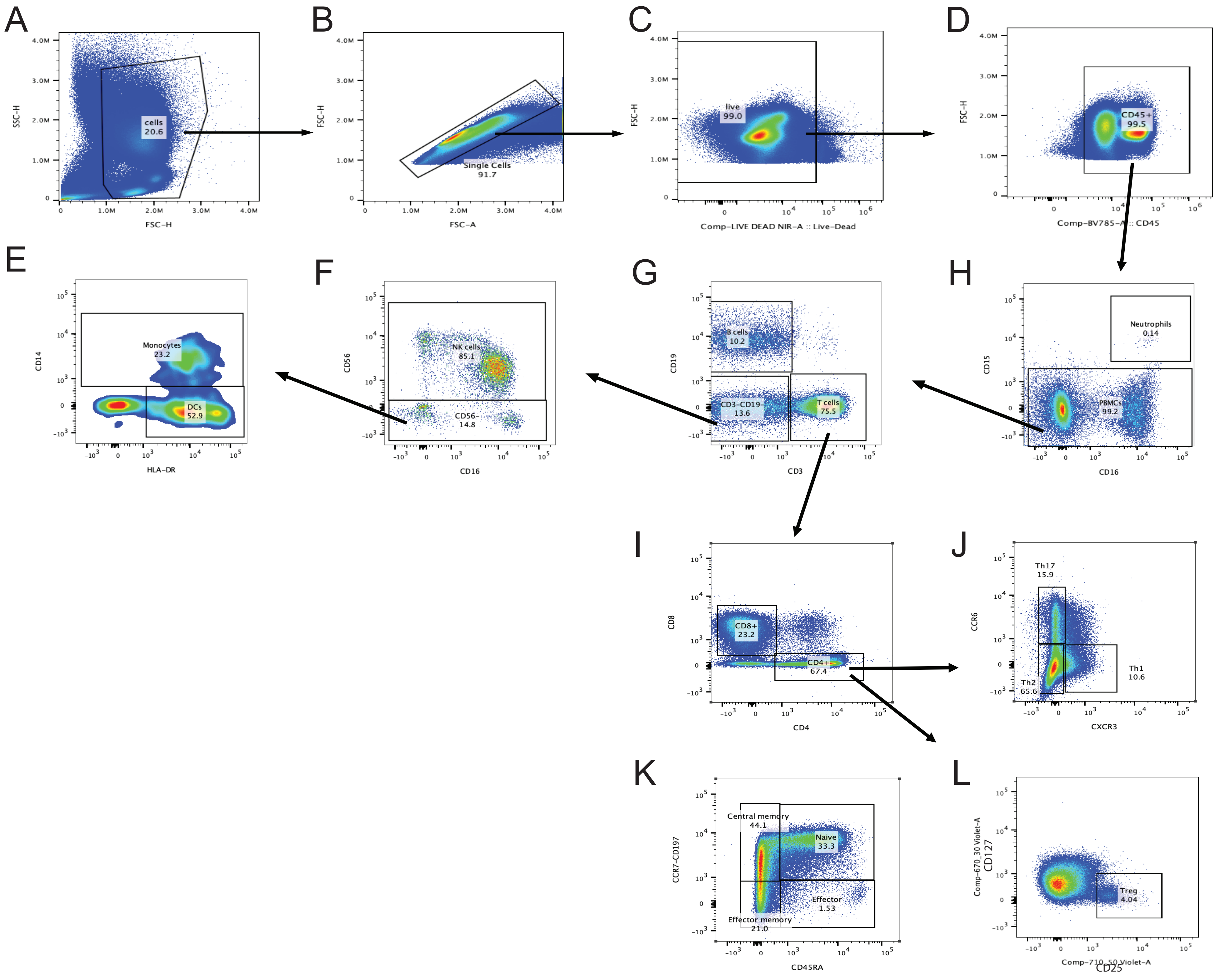

### Supplemental Figure 4

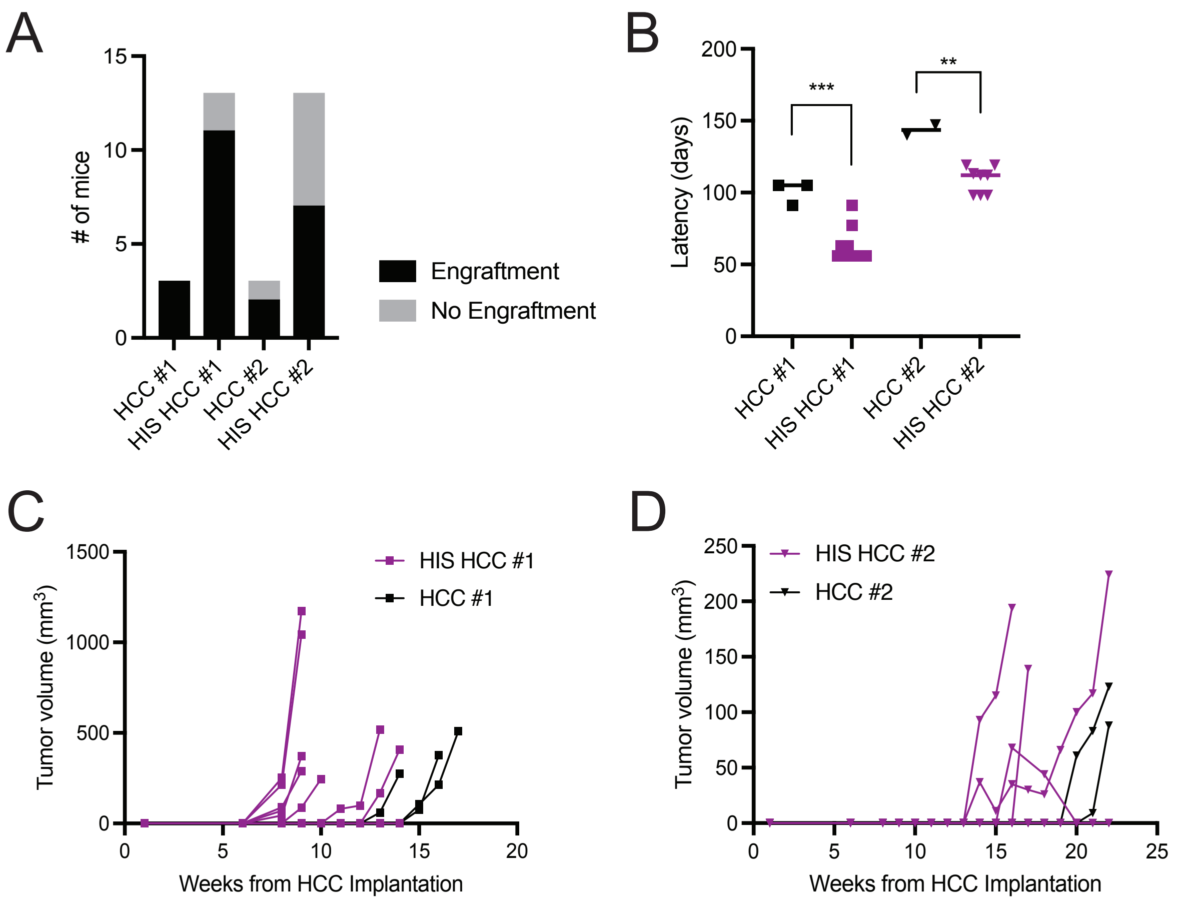
